# Functional selectivity for social interaction perception in the human superior temporal sulcus during natural viewing

**DOI:** 10.1101/2021.03.26.437258

**Authors:** Haemy Lee Masson, Leyla Isik

## Abstract

Recognizing others’ social interactions is a crucial human ability. Using simple stimuli, previous studies have shown that social interactions are selectively processed in the superior temporal sulcus (STS), but prior work with movies has suggested that social interactions are processed in the medial prefrontal cortex (mPFC), part of the theory of mind network. It remains unknown to what extent social interaction selectivity is observed in real world stimuli when controlling for other covarying perceptual and social information, such as faces, voices, and theory of mind. The current study utilizes a functional magnetic resonance imaging (fMRI) movie paradigm and advanced machine learning methods to uncover the brain mechanisms uniquely underlying naturalistic social interaction perception. We analyzed two publicly available fMRI datasets, collected while both male and female human participants (n = 17 and 18) watched two different commercial movies in the MRI scanner. By performing voxel-wise encoding and variance partitioning analyses, we found that broad social-affective features predict neural responses in social brain regions, including the STS and mPFC. However, only the STS showed robust and unique selectivity specifically to social interactions, independent from other covarying features. This selectivity was observed across two separate fMRI datasets. These findings suggest that naturalistic social interaction perception recruits dedicated neural circuity in the STS, separate from the theory of mind network, and is a critical dimension of human social understanding.

**Significance Statement:** Social interaction perception guides our daily behavior, yet it is unknown how our brain processes real-world social interaction scenes. Here, we demonstrate that social brain areas, including the superior temporal sulcus (STS) and medial prefrontal cortex (mPFC), are sensitive to broad social-affective information in naturalistic movies, replicating prior results with controlled paradigms. We show for the first time however, that the STS, but not mPFC, selectively processes social interactions in natural movies, independent of other co-occurring perceptual and social features, including motion, presence of faces, speech, and mentalizing about others. Our results suggest that social interaction perception is a crucial dimension of social understanding represented in the human brain.

## Introduction

Humans learn a great deal about the social world by observing others’ social interactions, defined here as action or communication between two or more people that are directed at and contingent upon each other. This social observation helps us form impressions of interacting people (e.g., whether they are prosocial or antisocial), their social relationships, and relative social status (Quadflieg and Koldewyn, 2017). Our ability to extract social information about interacting people appears to emerge in infancy (Hamlin and Wynn, 2011) and is preserved across species (Cheney and Seyfarth, 1986), underscoring the importance of this observational ability.

Previous studies have reported that the posterior superior temporal sulcus (STS) (Isik et al., 2017; Walbrin et al., 2018; Walbrin and Koldewyn, 2019) is selectively involved in recognizing others’ social interactions, in a manner that is functionally distinct from other related social functions, including detecting faces, animacy, other agents’ goals and theory of mind. These prior studies, however, have used simple stimuli, such as point-light figures or staged images that are largely devoid of other social information. It thus remains unknown how the human brain processes social interactions in real world scenes, where the interactions are highly confounded with other perceptual and social properties, including motion, faces, and theory of mind. The goal of the current study is to identify the brain mechanisms underlying real-world social interaction perception by adopting a naturalistic movie viewing paradigm.

A growing body of evidence in visual neuroscience suggests that naturalistic neuroimaging paradigms better uncover neural representations of objects, faces, scenes, and actions (Nishimoto et al., 2011; Wen et al., 2018; Haxby et al., 2020). Natural movies elicit stronger and more reliable brain responses than traditional experiments (Hasson et al., 2010; Sonkusare et al., 2019; Meer et al., 2020). In contrast, social neuroscience has not yet fully exploited these methods, although improving ecological validity in (social) cognitive neuroscience is considered an urgent challenge (Nastase et al., 2020; Redcay and Moraczewski, 2020). While several recent social neuroscience studies have adopted this paradigm to investigate theory of mind (Jacoby et al., 2016; Richardson et al., 2018; Richardson, 2019) and affective experience (Chen et al., 2020), only one study has investigated how the human brain processes social interactions during movie viewing (Wagner et al., 2016). This study revealed the involvement of the mPFC in social interaction perception, conjecturing that others’ social interactions invite a viewer to infer others’ personality and intention. However, it is unclear how the presence of other social features in the movie, particularly theory of mind, influenced these findings, which contribute to a longstanding debate about the distinct vs. overlapping roles of social perception and mentalization. Moreover, these results are in conflict with prior studies that used animated videos (Lahnakoski et al., 2012) and the above-mentioned studies with controlled stimuli, which identified the STS in social interaction perception. As a result, the neural underpinnings of social interaction perception, particularly in naturalistic contexts, remain unknown.

Here, we implemented computer vision techniques, machine-learning-based encoding model analyses, and variance partitioning to identify the unique contribution of social interactions to responses in the human brain. To improve ecological validity and replicability, we answered this novel question by applying the same analyses on two different fMRI datasets (Chen et al., 2017; Aliko et al., 2020) collected by different labs showing movies from different genres (crime vs. romance) to groups of participants living in different countries (US vs. UK) on different MRI scanners (3T vs. 1.5T). Our findings reveal that that social-affective features, consisting of an agent speaking, social interactions, theory of mind, perceived valence, and arousal, independently contribute to predicting brain responses in the STS and the mPFC. However, we find that the STS, but not mPFC, shows unique selectivity for others’ social interactions in particular, independent of other co-varying features.

## Materials and Methods

### fMRI data sources

We analyzed two publicly available fMRI datasets from two different studies. In the first study (Chen et al., 2017), 17 participants (male = 10) watched the first episode of the Sherlock BBC TV series (duration ~ 45 mins) in the scanner. In the second study (Aliko et al., 2020), 86 participants watched 10 different movies from 10 genres. We selected 20 participants who watched a commercial movie, 500 Days of Summer (duration ~ 90 mins). We excluded two participants from the second study as one participant (ID 14 in the original study) was scanned with a different head coil, and another participant (ID 16 in the original study) was offered glasses only after the first run, leaving 18 participants (male = 9) for our second movie analyses.

### fMRI data acquisition and preprocessing

In the first study (Sherlock), fMRI data were obtained on a 3T Siemens Skyra scanner with a 20-channel head coil. Whole-brain images were acquired (27 slices; voxel size = 4 × 3 × 3 mm^3^) with an echo-planar (EPI) T2*-weighted sequence with the following acquisition parameters: repetition time (TR) = 1500 ms, echo time (TE) = 28 ms, flip angle (FA) = 64°, field of view (FOV) = 192 × 192 mm^2^. In the second study (Summer), fMRI data were obtained on a 1.5T Siemens MAGNETOM Avanto with a 32-channel head coil. Whole-brain images were acquired (40 slices; voxel size = 3.2 × 3.2 × 3.2 mm^3^) with a multiband EPI sequence with the following acquisition parameters: multiband factor = 4, no in-plane acceleration, TR = 1000 ms, TE = 54.8 ms, FA = 75°.

We obtained preprocessed fMRI data from the authors of each original study. In the original Sherlock study, preprocessing steps included slice-timing correction, motion correction, linear detrending, temporal high-pass filtering (140s cut off), spatial normalization to a Montreal Neurological Institute (MNI) space with a re-sampling size of 3 × 3 × 3 mm^3^, and spatial smoothing with a 6-mm full width at half maximum (FWHM) Gaussian kernel. fMRI data were also shifted by 4.5 s (3 TRs) from the stimulus onset to account for the hemodynamic delay. Lastly, timeseries blood oxygenation level dependent (BOLD) signals were z-score standardized.

In the Summer study, authors performed slice-timing correction, despiking, motion correction, spatial normalization to MNI space with a re-sampling size of 3 × 3 × 3 mm^3^, and spatial smoothing with a 6-mm FWHM Gaussian kernel. Timeseries BOLD signals were scaled to 0~1 and detrended based on run lengths varying across participants and runs, head-motion parameters, and averaged BOLD signals in white matter and cerebrospinal fluid regions. Timing correction was also applied to align the fMRI timeseries and the movie. Detailed information on how movie was paused and restarted for each run and timing correction related this issue can be found in the original study (Aliko et al., 2020). In addition, similar to the Sherlock study, we shifted Summer fMRI by 4 s (4 TRs) from the stimulus onset to account for the hemodynamic delay.

### Movie analysis and annotations

To examine how BOLD responses relate to each stimulus feature, we fit a linear regression model where voxel-wise responses are predicted based on a linear combination of stimulus features (Fig 1A). To create a stimulus feature space, we annotated the movies with a mix of fully automatized approaches and human labeling. We first split the full-length Sherlock episode and the Summer movie into 1.5 and 3 sec segments, respectively. We excluded the introductory video clip, a short-animated movie shown before the Sherlock episode in the original study, from the analysis. For the Summer movie, the opening and ending credits were excluded from the analysis. fMRI volumes matching these scenes were truncated and excluded from further analysis. A total of 1921 and 1722 video segments were generated from the Sherlock and Summer stimuli, respectively.

**Fig 1A.**
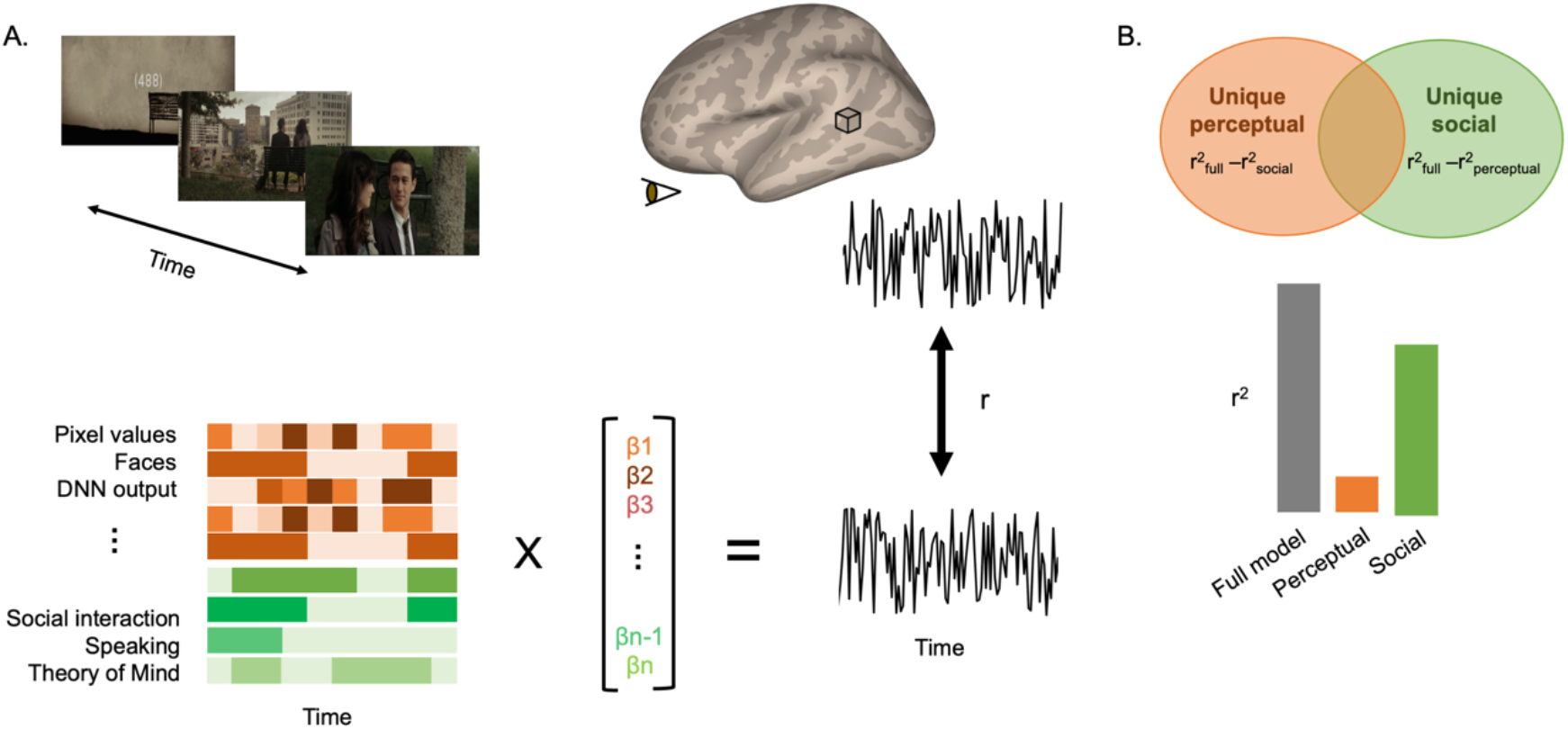
Encoding model overview. We labeled perceptual and social-affective features from a movie that participants viewed in the MRI scanner. The features include hue, saturation, value for each pixel, motion energy, the presence/absence of written words, indoor vs. outdoor scenes, the presence/absence of faces, audio amplitude, pitch, the presence/absence of music, the presence/absence of social interactions, the presence/absence of an agent speaking, the presence/absence of agent talking about others’ mental states (theory of mind), perceived valence, and arousal of the scene for both movies, and additionally the social touch feature for the Summer movie. During model training, we learned a set of beta weights linking the features to fMRI BOLD responses over time. We then predicted the response to held-out movie data by multiplying the movie feature vectors by their corresponding beta weights, and correlated these predictions with the actual responses extracted while participants viewed the movie. As a result, a prediction performance score (r) of the model is assigned to each voxel. B. Variance partitioning analysis overview. Prediction performance r values from the full, perceptual, and social-affective models are used to calculate unique variance explained by the perceptual or social-affective model for each voxel.

Using MATLAB (R2020a, The Mathworks, Natick, MA) built-in functions, we extracted low-level visual features, namely hue, saturation, pixel values (HSV) and motion energy, and auditory features, namely amplitude and pitch. Specifically, we computed HSV (‘rgb2hsv’) and motion energy (‘opticalFlowFarneback’) in each pixel for all frames (640 × 360 pixels and 38 frames for Sherlock, 720 × 576 pixels and 75 frames for Summer) and averaged these values over pixels and frames. In both movies, the majority of motion comes from humans moving so we believe the motion energy to also be a good proxy for biological motion. We also extracted high-level visual features from a deep convolutional neural network model (DNN, see below section). For audio amplitude (‘audioread’) and pitch (‘pitch’), we averaged values over the audio samples and channels (66150 samples and two channels for Sherlock and 132300 samples and two channels for Summer). This computation allowed us to obtain one value per feature for each video segment (the duration of 1 TR).

The authors of the Summer study (Aliko et al., 2020) used the ‘Amazon Rekognition’ service (https://aws.amazon.com/rekognition/) to obtain faces annotations. Based on their annotations, we extracted the presence or absence of faces in each segment. We implemented the same analysis pipeline (‘start_face_detection’ and ‘get_face_detection’ functions from the Amazon Rekognition) to the Sherlock video segments. We used the binary label (presence of a face(s) = 1, absence of a face = 0) for each video segment with the confidence level 99% as a threshold.

We also included some of the publicly available annotations, made by human raters in the original Sherlock study (Chen et al., 2017) – whether the location of the scene is indoor or outdoor (indoor = 1, outdoor = 0), whether or not music plays in the background (presence of music = 1, absence of music = 0), and whether or not there are written words on the screen (presence of written words = 1, absence of written words = 0). In the current study, two human raters made the same annotations for the Summer video segments.

Two human raters indicated whether or not a video segment contains social interactions (defined as actions or communication between two or more individuals that are directed at and contingent upon each other, presence of social interactions = 1, absence = 0). These labels were highly consistent across our two raters (r = 0.92 and r = 0.86 for Sherlock and Summer, respectively). The raters also rated additional social features, including whether or not a person speaks in the scene (yes = 1, no = 0), and whether or not a person infers mental states of others (yes = 1, no = 0). The annotation of the theory of mind feature is based on whether a movie character is inferring other characters’ thoughts and emotions in each scene (an example in Sherlock – Ms. Patterson is seated at a table at a press conference, reading her statement saying: “He loved his family and his work”; an example in Summer – The main character seen sitting next to other colleagues at a meeting and a narrator says: “The boy, Tom Hansen of Margate of New Jersey, grew up believing that he’d never truly be happy until the day he met the one.”). We selected this criterion to be as objective as possible, and to avoid raters guessing about whether the subjects were engaged in mentalization. This type of second-order theory of mind task activates the theory of mind network (Tholen et al., 2020). As in prior studies (Saxe and Powell, 2006), the description of a character’s appearance (e.g., she is tall and thin) or bodily sensation (e.g., she had been sick for three days) were not considered theory of mind. For Summer, we additionally included a touch feature – whether or not a person makes physical contact with another person (social touch = 1), him(her)self or an object (nonsocial touch = −1), or not (absence of touch = 0). We were unable to include the touch feature in the Sherlock data as there were less than 10 video segments displaying social touch.

Considering that the time-scale of high-level cognitive events is longer than a low-level perceptual event (Baldassano et al., 2017), we merged each pair of consecutive Sherlock 1.5s length segments into one for annotations of social features, resulting in 3s length segments for both studies. For binary features, the relative frequency of feature occurrence is described in Table 1.

**Table 1.** Relative frequency of feature occurrence in each movie. Only binary features are included. Note that multiple annotators used binary scores to rate these features, and their ratings were averaged, yielding values ranging from 0 to 1. In the case of decimals, we round them to the nearest whole number (0 or 1) when counting the occurrence.

|  | Written Text | Music | Indoor | Face | Speaking | Social Interaction | Theory of Mind | Touch |
| --- | --- | --- | --- | --- | --- | --- | --- | --- |
| Sherlock | 4% | 63% | 71% | 85% | 65% | 75% | 32% | not included |
| Summer | 19% | 66% | 66% | 92% | 58% | 77% | 20% | 11% |

Lastly, we added valence and arousal ratings, offered by the authors of another Sherlock fMRI study (Kim et al., 2020). In their study, 4.5s video segments, parsed from the full-length Sherlock episode, were rated by 113 participants with a 1 – 9 Likert scale. 35 participants rated all video segments, and the remaining participants rated only a quarter, yielding about 55 ratings per video segment. Group mean valence and arousal ratings were used in the current study. For Summer segments, four human raters judged how pleasant and arousing the scene is using a 1 – 5 Likert scale. Despite the small number of raters, inter-rater consistencies are relatively high (valence Spearman r (r_s_) = 0.62, arousal r_s_ = 0.35), compared to that (valence r = 0.30) reported in the Sherlock study (Kim et al., 2020).

In summary, perceptual features consist of components from the DNN fifth layer (see below), HSV, motion energy, faces, indoor/outdoor, written words, amplitude, pitch, and music. Social-affective features consist of social interactions, speaking, theory of mind, valence, and arousal for both experiments, and additionally the touch feature for the Summer data. Fig 2 illustrates correlations across one-dimensional features. Although these feature spaces are moderately correlated, we believe them to capture important non-overlapping information. We ran principal component analysis (PCA) on the five and six social-affective features labeled in the Sherlock and Summer movies, respectively. According to this analysis five (Sherlock) and six (Summer) principal components are needed to account for 99% of the feature variance, suggesting that the dimensionality of these feature spaces cannot be reduced without losing explanatory power.

**Fig 2.**
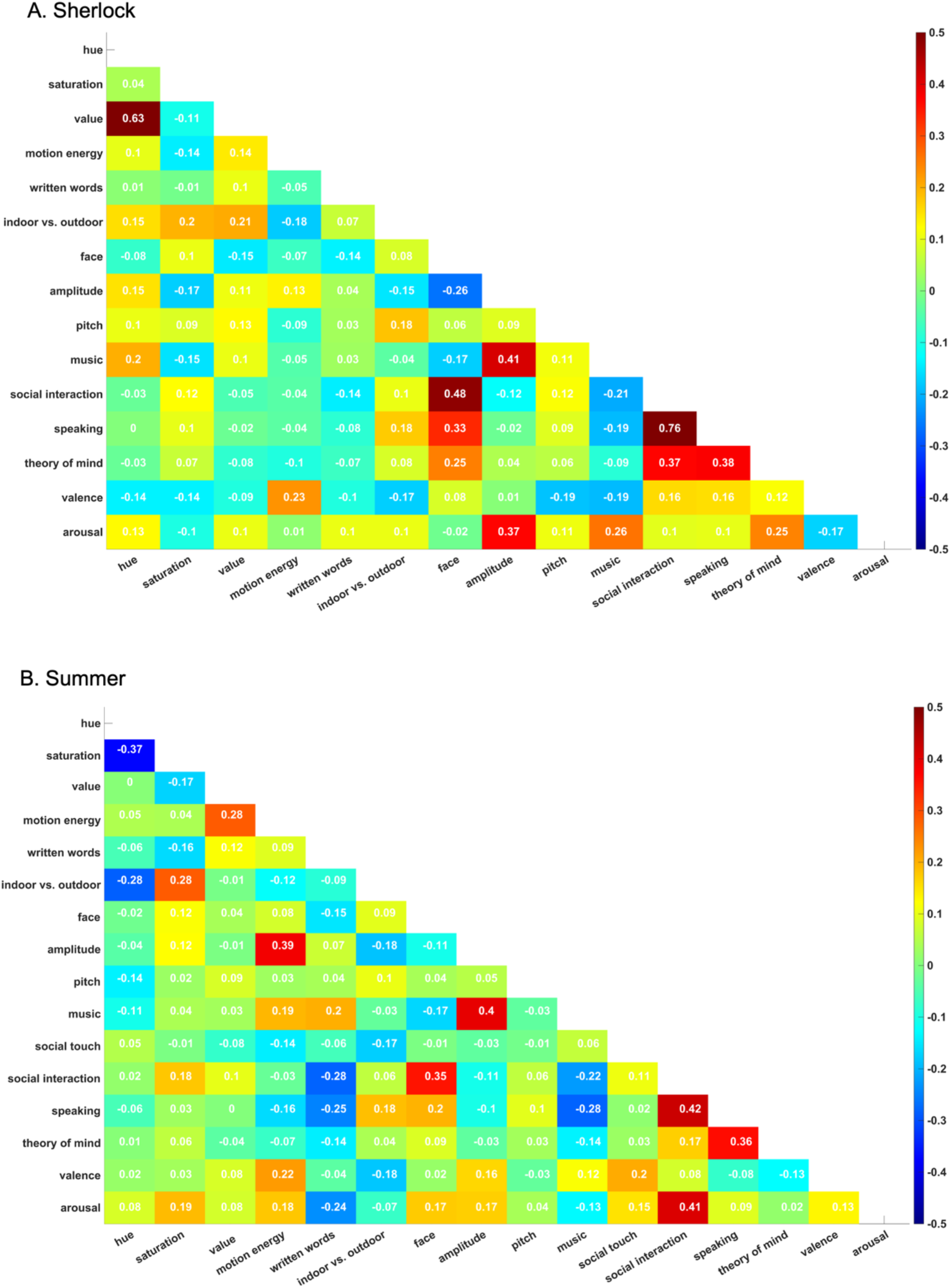
The pairwise rank correlations between features in Sherlock (A) and 500 days of Summer (B) movie. Cells in the matrix show the correlation coefficients between features, indicated by Spearman r values and color coding (red: positive correlation; blue: negative correlation). DNN features are not included in this correlational analysis as they are high-dimensional, unlike other features (1 vector per feature).

### Deep convolutional neural network

We used a deep convolutional neural network to extract visual features from both stimuli. We included DNN features as they have recently been shown to explain a great deal of variance throughout both early and later stages of visual cortex (Wen et al., 2018). Specifically, PyTorch (version 1.4.0) implementation of AlexNet (Krizhevsky et al., 2012), pretrained on the ImageNet dataset (Russakovsky et al., 2015), was used. AlexNet consists of eight layers with five convolutional layers and three fully connected layers. AlexNet in PyTorch adopted 64, 192, 384, 256, and 256 kernels from the first through fifth layers instead of 96, 256, 384, 384, and 256 reported in the original study (Krizhevsky et al., 2012). The size of the kernels is the same as original AlexNet – 11 × 11, 5 × 5, 3 × 3, 3 × 3, 3 × 3 from the 1^st^ to 5^th^ layers, respectively.

The first frame of each video segment was, cropped, normalized, and fed into the first layer of AlexNet, with the image input size 3 × 224 × 224. The first frame was chosen as it matches the onset of each TR and visual scenes do not change dramatically every 1.5 or 3s in the movies. The output of the previous layer served as the input of the following layer. Including all DNN layers in the encoding model generates more features than samples, which risks overfitting, and is computationally expensive. We captured visual features of each video segment by taking the activations of all units from the fifth layer right before the rectified linear activation function and the max-pooling layer during the forward pass. As activations from the fifth layer are highly correlated to those of early (r > 0.75 with second layer) and late layers (r > 0.5 seventh layer), we assume that the fifth layer captures low, mid, and high-level visual information shown in the movies without risking overfitting. The output size of the fifth layer is 256 × 13 × 13, which was flattened to an array with 43264 elements. Unlike other features (1 × the number of video segments), the features from the DNN layer are high dimensional (i.e., 43264 × the number of video segments). We reduced the dimensionality without losing crucial information by applying PCA to the DNN activations. Components first through N^th^ were selected until the amount of explained variance reached 70%. 147 and 150 components were added to the feature matrix of the fifth layer for Sherlock and Summer segments, respectively.

### Inter-subject brain correlation

Inter-subject correlation (ISC) analysis was implemented to create a brain mask containing voxels showing shared stimuli-evoked responses across participants. ISC is a well-validated fMRI method that identifies voxels with reliable neural responses across time while rejecting idiosyncratic and noisy voxels (Nastase et al., 2019). Using the brain imaging analysis kit’s (BrainIAK, version 0.1.0) built-in functions (‘isc’, ‘permutation_isc’), we measured ISC with a leave-one-subject-out approach. In other words, for every voxel, time-series BOLD responses for all but one subject were averaged and correlated with that of the remaining subject. We repeated this process for every participant. Fisher-Z transformed correlation values were averaged across folds. Encoding model analysis was masked to include only voxels with ISC value > 0.25 in the case of the Sherlock dataset, based on the previous permutation results (P _the false discovery rate (FDR)_ < 0.001) (Baldassano et al., 2017). For the Summer dataset, we ran a sign permutation test (5,000 iterations) across participants, as implemented in BrainIAK, and created a mask consisting of voxels passing the statistical threshold, P _FDR_ < 0.005 after the multiple comparison correction with the FDR. In other words, subjects’ ISC values were randomly multiplied with +1 or −1 for 5,000 times, which resulted in empirical null distribution of ISC values for each voxel. From this distribution, two-tailed P values were calculated and adjusted with FDR correction. We used a less stringent threshold (P _FDR_ < 0.005 as opposed to P _FDR_ < 0.001) on Summer ISC to have a mask with the similar number of voxels with that of Sherlock (25468 and 22044 voxels, respectively). Voxels outside of the mask or whose BOLD responses did not change over time were excluded from the further encoding analyses.

### Voxel-wise encoding modeling

We used an encoding model approach to predict the voxel-wise BOLD responses evoked during natural movie viewing. For each voxel, BOLD responses were modeled as a linear combination of various feature spaces (Fig 1A). Each feature was normalized over the course of the movie. Specifically, banded ridge regression was implemented to estimate the beta weights of stimulus features in the nested cross-validation scheme. Unlike classical L2-regularized ridge regression, banded ridge regression does not assume all feature spaces require the same level of regularization and allows more than one ridge penalty in the prediction model (Nunez-Elizalde et al., 2019). Thus, banded ridge regression has advantages over ordinary least squares regression and classical ridge regression. It minimizes overfitting to noise during training, and it is a preferred method when the feature spaces are high-dimensional or suffer from (super)collinearity, ultimately improving the prediction performance (Nunez-Elizalde et al., 2019). In particular, we used two different ridge penalties: one for the high-dimensional DNN features and a second for all other single-dimensional (per video segment) features. We did this to avoid an overweighting of the DNN features that may occur with one shared ridge penalty.

Data were first split into 10 folds. Among them, nine folds were used to estimate beta weights and optimize the regularization parameter (range 0.1 ~ 10,000) λ_1_ and λ_2_. Optimal λ values were selected per voxel via inner loop 5-fold cross-validation. In other words, training data from nine folds were again split into five folds. Four folds were used for the model estimation with various λ values, and unseen data from the remaining fold in the inner loop were used for selecting the λ values that, on average, yielded the smallest mismatch between predicted and actual BOLD responses. After the model estimation and regularization parameter optimization, unseen data from the remaining tenth fold in the outer loop were used for evaluating the performance of the model. We measured model performance by computing the correlation between predicted and actual BOLD responses. For each voxel separately, this process was repeated 10 times, and the performance of the model was averaged over the repetitions.

Specifically, our encoding model was built as follows. The first step is to normalize the dependent (Y) and independent (X) variables by centering and scaling them to have mean 0 and standard deviation 1. Let Y be an array of size T_r_ (number of total fMRI volumes from the training set) consisting of the zero-centered BOLD signal amplitudes after the normalization. Let X_1_ and X_2_ be a T_r_ × F_1_ (number of DNN features) and T_r_ × F_2_ (number of additional features) matrix, respectively. Y in the banded ridge regression is then:

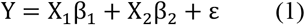

where β_1_ and β_2_ are a F_1_ or F_2_ sized array containing the beta weights for each feature.

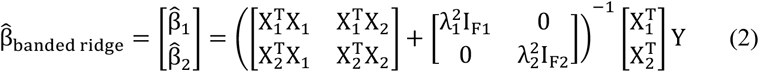

Note that when λ_1_ = λ_2_, a regression model becomes classical ridge regression, and when λ_1_,λ_2_ are 0, a model becomes ordinary least squares regression. Having an option of λ_1_ = λ_2_ or λ_1_ = λ_2_ = 0, banded ridge regression performs at least as good as classical ridge or ordinary least squares regression. The size of the diagonal matrix containing λ_1_ and λ_2_ is F × F, where F = [F_1_ F_2_]. The diagonal entries consist of repeats of λ_1_, as many as the size of array F_1_, and repeats of λ_2_ for the remaining entries. As described earlier, the optimal λ values were selected in the inner loop.

Using estimated beta weights, 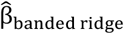, unseen BOLD responses were predicted as:

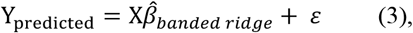

where Y_predicted_ is an array of size T_e_ (number of total fMRI volumes from the testing set) consisting of estimated BOLD signals. Let X = [X_1_ X_2_], where X_1_ and X_2_ are a T_e_ × F_1_ and T_e_ × F_2_ matrix, respectively. The last step is to calculate correlation coefficient value between the predicted and actual BOLD signal to evaluate the model performance. A high r value can be interpreted as high prediction performance.

Three separate encoding models were built to evaluate which voxel could be accurately predicted by the linear combination of all, perceptual, or social-affective features.

1. A full model includes all features listed in the sub-section (movie analysis and annotations). In this model, X_1_ is composed of high-dimensional DNN components and X_2_ is composed of the rest. Beta weights estimated from this model were used in preference mapping analysis described below.
2. A perceptual model includes all visual and auditory features. X_1_ is composed of DNN components and X_2_ is composed of the rest of visual and auditory features.
3. A social-affective model includes all social-affective features. In the social-affective model, the feature space is low-dimensional, so classic ridge regression (where λ_1_ = λ_2_) was used. Thus, X is composed of all social-affective features.

We calculated the prediction performance map across participants for every model. Random-effect group-level analyses were conducted with a nonparametric permutation test to identify voxels showing significantly above chance prediction performance. Before the permutation test, we removed voxels that were not shared across participants, which resulted in the final number of voxels being slightly less than the total number of voxels contained in the mask (21649 voxels for Sherlock; 24477 voxels for Summer). Similar to previous studies (Cichy et al., 2017; Hebart et al., 2018), and the above-mentioned ISC analysis, we conducted a sign permutation test (5,000 iterations). From the empirical null distribution of a prediction performance, one-tailed P values were calculated and adjusted with FDR correction. Group averaged prediction performance maps of each model were thresholded at P _FDR_ < 0.05 and plotted on the cortical surface. Note that we report original r values as no noise ceiling (upper limit of model performance) normalization was performed.

Two additional encoding models were built to evaluate the unique effects of two main social features of interest, the presence of a social interaction and mentalization, on the BOLD response prediction.

4) One model includes all features except the presence of a social interaction
5) The final model includes all features except the mentalization feature.

Prediction performances obtained from these five models were used in variance partitioning analyses. All code for voxel-wise encoding and variance partitioning analysis along with new movie annotations are available at https://github.com/haemyleemasson/voxelwise_encoding.

### Preference Mapping

To gain a comprehensive understanding of the relative contribution of different features to activity in each voxel, we performed preference mapping analysis to visualize the stimulus features that were best and the next best at explaining each voxel’s activation. The primary advantage of this method is that it allows us to consider all features in a single analysis. Beta weights generated from a full model were used to predict the withheld BOLD responses for each voxel via a cross-validation as described above. During the testing session, we used the beta weight(s) of a feature of interest (e.g., the amplitude of audio) to estimate unseen BOLD responses while assigning 0 values to beta weights of other features. This method is superior to having a separate model for each feature when evaluating the relative contribution of each individual feature in predicting voxel activations (Nunez-Elizalde et al., 2019). Moreover, this method is suitable for examining the prediction performance for high-dimensional feature spaces, such as DNN units, which produces hundreds of beta weights. We repeated this procedure for every feature and participant. In the end, the winning feature that yielded the highest group averaged prediction performance and the next highest were selected for each voxel. We colored every voxel to reflect which feature predicts BOLD responses of that voxel the best and the second best. The first and second preference maps were plotted on the cortical surface.

### Variance Partitioning

While a preference mapping analysis measures the relative contribution of each stimulus feature in predicting held out BOLD responses, a variance partitioning analysis determines the unique contribution of each feature or group of features to this prediction. Unlike preference mapping it is not feasible to compare all features individually in this analysis, thus we used this to better understand the unique contributions of two broad groupings of features (perceptual and social-affective) and two important social features (the presence of a social interaction and theory of mind). To this end, we compared the amount of variance explained by a perceptual or social-affective model with that of a full model consisting of all features (Fig 1B), as well as the unique contribution of the presence of a social interaction and mentalization, in predicting the BOLD responses. The amount of unique variance explained by a model/feature of interest was calculated as:

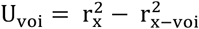

X reflects all features listed in the sub-section (movie analysis and annotations). VOI reflects a variable of interest (e.g., all social-affective features or a mentalization feature). r^2^ is the squared prediction performance value obtained from the encoding model, which can be interpreted as the variance explained by a model consisting of either all features X or all features except the variable of interest X – VOI. U_voi_ is the amount of unique variance explained by VOI. We computed U_voi_ for every participant and voxel. Thus, U_voi_ is an N × V sized matrix where N reflects the total number of participants and V reflects the total number of voxels included in the encoding model analysis for each dataset (i.e., N=17 and Voxels = 21649 for Sherlock; N=18 and Voxels = 24477 for Summer). The higher the U value is, the more the variance is uniquely explained by a variable of interest. Like voxel-wise encoding, we applied the sign permutation test (5,000 iterations) for the statistical inference. Group averaged maps, resulting from U values, were thresholded at P _FDR_ < 0.05 and plotted on the cortical surface. Most brain areas were labeled with automated anatomical labeling atlas (Tzourio-Mazoyer et al., 2002). The location of STS and temporoparietal junction (TPJ) were compared with the templates created from the previous studies (Mars et al., 2012; Deen et al., 2015).

## Results

### Perceptual and social-affective features accurately predict voxel-wise activation throughout the brain

Two sets of subjects (N = 17 and N = 18) viewed two different movies (the first episode of the Sherlock BBC TV series and 500 Days of Summer) while their BOLD responses were recorded in fMRI. We extracted a range of perceptual and social features from each movie using a combination of automatic methods and human labeling. The perceptual features consisted of low-level sensory features including HSV, motion energy, audio amplitude and pitch, as well as higher-level perceptual features including the presence of faces, indoor vs. outdoor scenes, written words, the presence of music and features from the fifth (final) convolutional layer of DNN (see Movie analysis and annotations in Materials and Methods). Social-affective features consisted of social interactions, an agent speaking, theory of mind, perceived valence, and arousal for both studies, and additionally the touch feature for the Summer data. We performed voxel-wise encoding analyses to learn the relationship between these features and fMRI movie data, and then predicted held out BOLD responses using the features and beta values learned in training (Fig 1A). We focused our analyses on voxels with shared stimuli-evoked responses across participants as measured by ISC (see Inter-subject brain correlation in Materials and Methods).

In both fMRI datasets, the performance of the full model, consisting of all perceptual and social-affective features, was significantly better than chance at predicting voxel-wise responses throughout the brain. The full model significantly predicted BOLD responses in 100% and 99.99% of voxels inside of the ISC mask for the Sherlock (range of prediction performance (r) = 0.08 ~ 0.51, mean performance = 0.25, standard deviation (std) = 0.07) and Summer fMRI data (range = 0 ~ 0.43, mean = 0.17, std = 0.06), respectively. Note that prediction performances differ across these voxels as reflected in Figure 3. The highest model performance was observed in the left STS in both studies (Peak MNI coordinates X, Y, Z = −63, −24, −3 for Sherlock; X, Y, Z = −66, −18, 1 for Summer). High prediction performance was also found in the right STS (highest performance in right STS = 0.48 for Sherlock (X, Y, Z = 51, - 33, 0); 0.43 for Summer (X, Y, Z = 67, −21, −3)).

**Fig 3.**
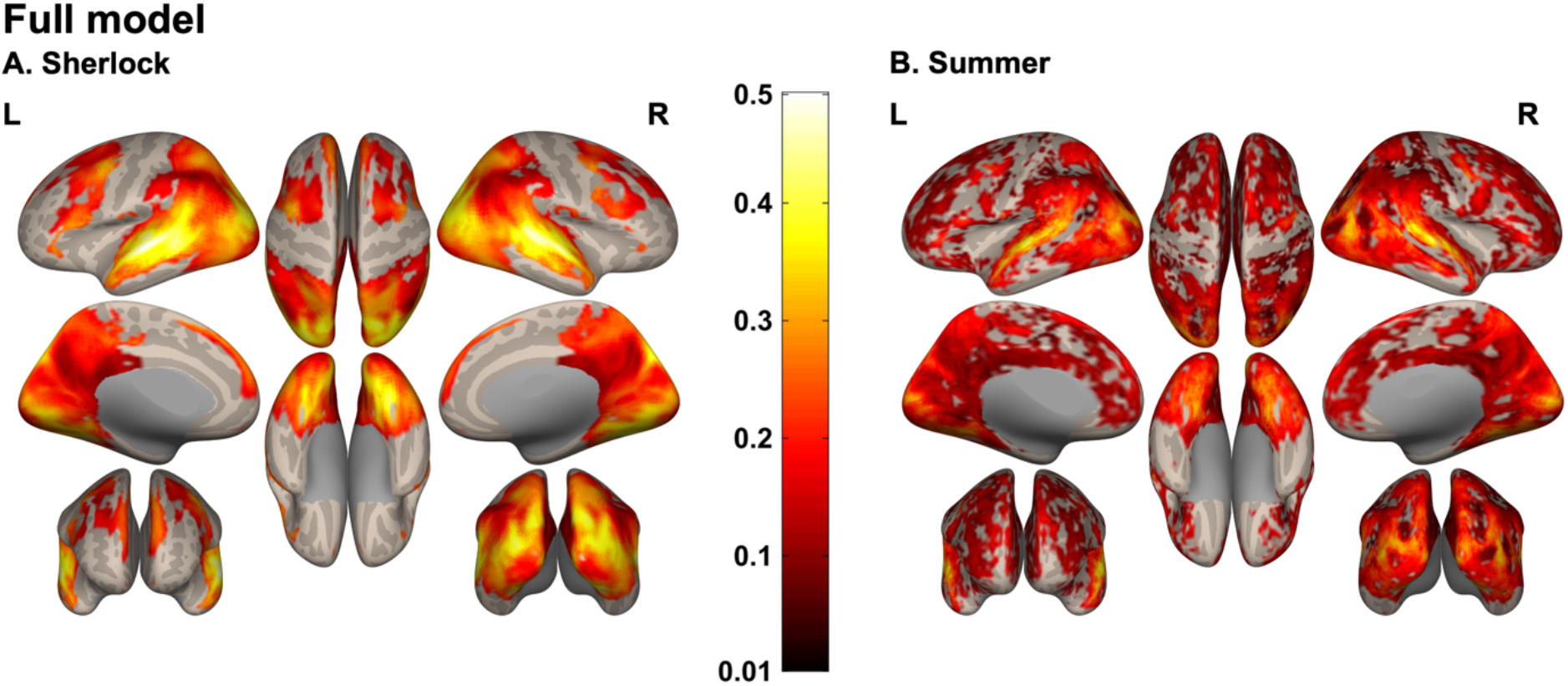
The full model prediction performance observed in Sherlock (A) and Summer (B) data. Group averaged performance scores are mapped on inflated cortices using the CONN software (Whitfield-Gabrieli and Nieto-Castanon, 2012) (P _FDR_ < 0.05, minimum cluster size >10 voxels). The color bar indicates the prediction performance score (0 = chance, 1 = perfect prediction). The full model predicts neural responses in the bilateral STS particularly well in both studies. L = left hemisphere, R = right hemisphere, FDR = the false discovery rate.

The perceptual model, consisting of the visual and auditory features listed above, significantly explained BOLD responses in 100% and 99.99% of voxels inside of the ISC mask in the Sherlock (range of prediction performance = 0.05 ~ 0.38, mean = 0.21, std = 0.06) and Summer data (range = 0 ~ 0.35, mean = 0.14, std = 0.05), respectively. The highest model performance was observed in the visual cortex in both experiments – the right fusiform gyrus (X, Y, Z = 27, −69, −12) in Sherlock and the early visual cortex (X, Y, Z = 13, −87, 7) in Summer (Fig 4).

**Fig 4.**
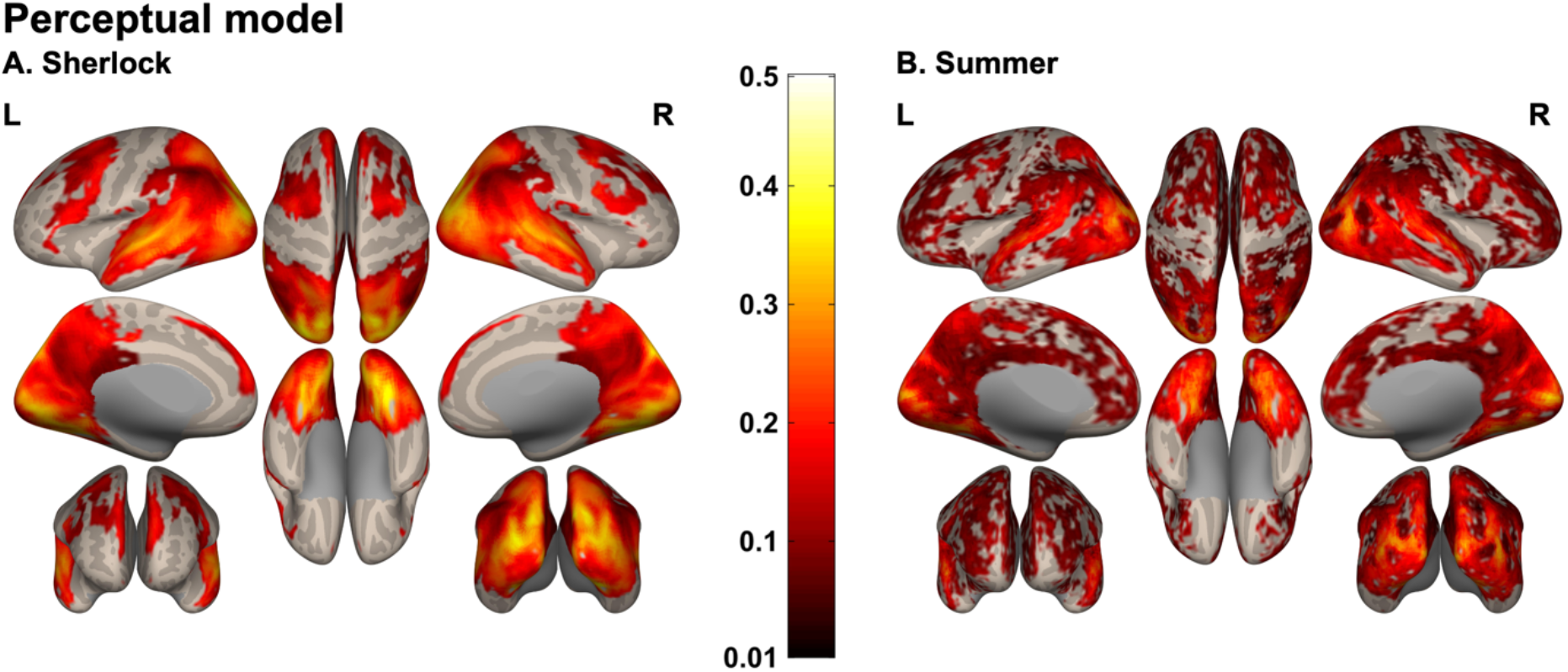
The perceptual model prediction performance observed in Sherlock (A) and Summer (B) data. The perceptual model significantly predicted neural responses in the visual cortex in both studies. Group averaged performance maps are thresholded at P FDR < 0.05 and the minimum cluster size >10 voxels.

A social-affective model, consisting of the social-affective features listed above (an agent speaking, social interactions, theory of mind, perceived valence, and arousal) also produced significantly above chance performance throughout the whole brain in both studies. The social-affective model explained significant variance in 100% and 99.92% of voxels inside of the ISC mask in the Sherlock (range of prediction performance = 0.04 ~ 0.55, mean = 0.20, std = 0.08) and Summer data (range = 0 ~ 0.39, mean = 0.12, std = 0.05), respectively. The highest model performance was observed in the left STS in both experiments (X, Y, Z = −60, −24, −3 for Sherlock; X, Y, Z = −63, −18, 1 for Summer). Again, findings are bilateral (highest performance = 0.49 in the right STS (X, Y, Z = 51, −33, 0) for Sherlock; 0.38 for Summer (X, Y, Z = 64, −24, 1)) (Fig 5).

**Fig 5.**
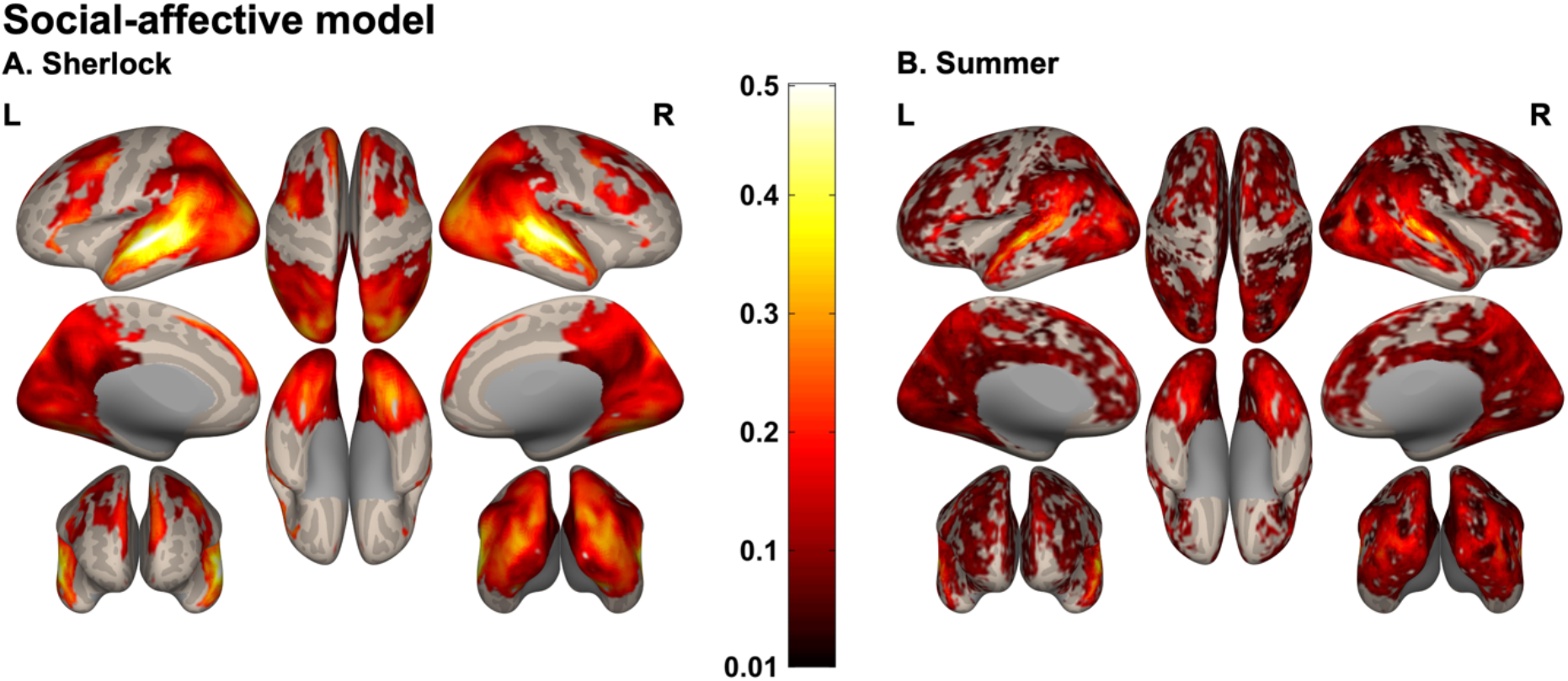
The social-affective model prediction performance observed in Sherlock (A) and Summer data (B). Group averaged performance scores are plotted on inflated cortices (P _FDR_ < 0.05, minimum cluster size >10 voxels). Like the full model, the social-affective model significantly predicted neural responses in the bilateral STS in both experiments.

### Social interaction features are the strongest predictor of STS activity, while deep neural network features are the strongest nearly everywhere else

We next performed preference mapping analyses to measure the relative contribution of each stimulus feature in predicting held out BOLD responses for every voxel. Fig 6 illustrates winning features that best (or second best) capture voxel-wise responses in each movie. The DNN fifth layer was the most predictive feature for most voxels throughout the brain (pink in Fig 6A-B). Notably, however, this was not true in the bilateral STS where two social interaction-related features, an agent speaking (green in Fig 6) and presence of social interactions (green-yellow in Fig 6), are the most or second most preferred stimulus features in both studies. As expected, the auditory cortex was best explained by the audio amplitude feature (yellow-orange in Fig 6) in both studies. Visual features, i.e., motion energy, faces, and written words, are the second-best features explaining neural responses in ventral and dorsal visual pathways (red colors in Fig 6C-D). TPJ and mPFC were second-best explained by social-affective features (theory of mind, speaking, social interaction, and arousal) (green and purple colors in Fig 6C-D). Frequency of feature occurrence does not seem to predict voxel preference. Although faces, indoor scenes, and background music are frequently present in both movies (Table 1), these features are not the ones preferred by most voxels.

**Fig 6.**
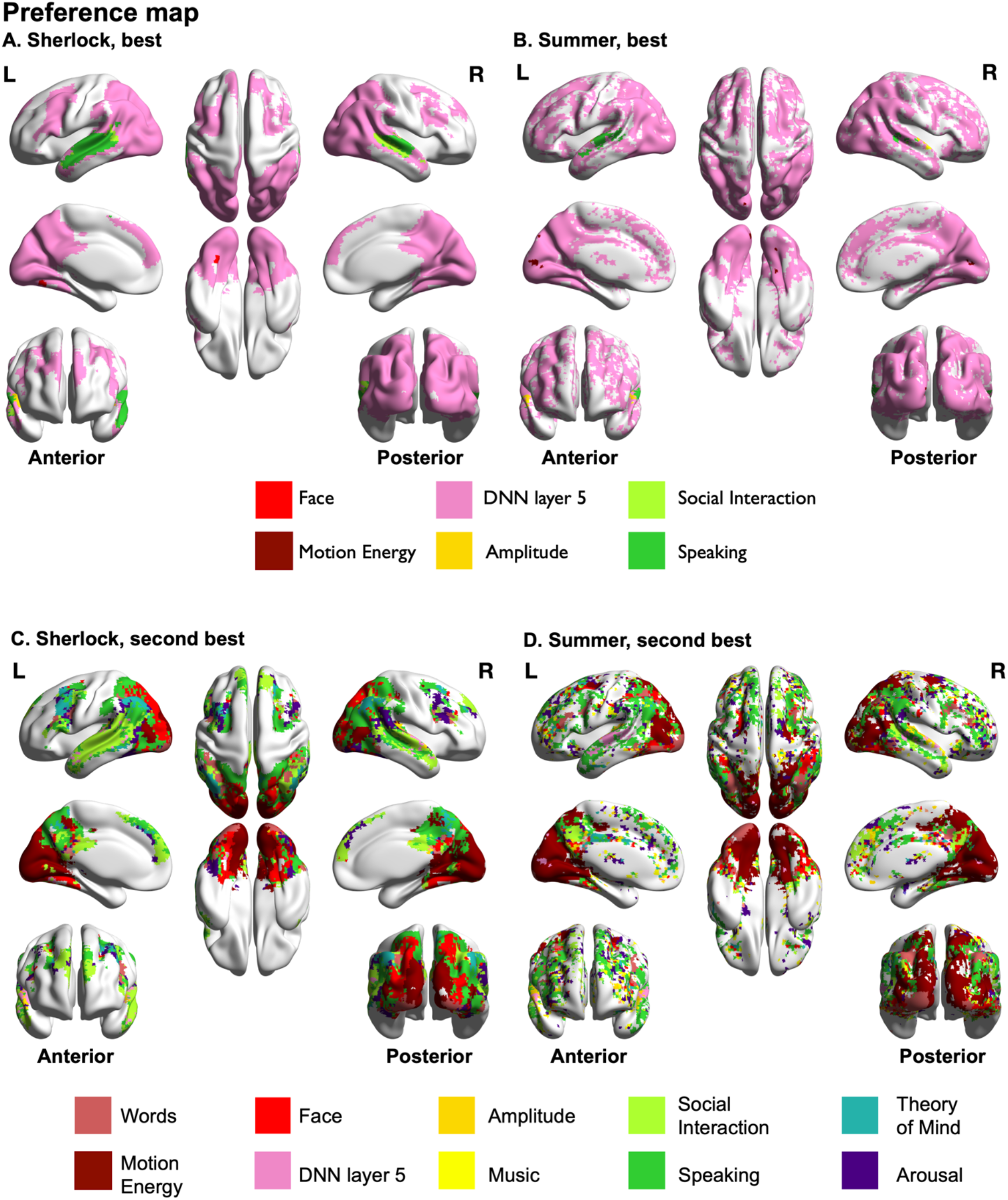
Preference maps for the best (panel A and B) and second-best features (panel C and D) in each voxel for Sherlock (A and C) and Summer (B and D) data. Feature color codes indicated at bottom. DNN features best explain the neural responses throughout most of the brain in both studies, except the STS, whose neural responses are best explained by social features. Overall, social-affective features (green and purple colors) are the most or at least the second most preferred in the temporal and frontal lobe in both studies, whereas visual features (red colors), in addition to DNN features, are preferred in the visual cortex.

### The perceptual model uniquely predicts responses in the visual and auditory cortex, while the social-affective model does so in the temporal and frontal regions

While above results indicate that STS and nearby regions in the temporal lobe are best explained by social features, they do not reveal to what extent perceptual and social-affective features contribute to the neural response, independently of their co-varying perceptual features. By conducting variance partitioning analyses (Fig 1B), we examined the unique contribution of the perceptual and social-affective feature models to the prediction of BOLD responses throughout the brain. This analysis is particularly important as many features are at least somewhat correlated with each other (e.g., rank correlation between social interactions and the presence of faces r = 0.48 in Sherlock and r = 0.35 in Summer, Fig 2).

The results indicated that the perceptual model significantly explained unique variance in brain regions implicated in visual or auditory processing (Fig 7A-B). The largest portion of unique variance explained was found in the early visual cortex in both studies (X, Y, Z = −6, −90, 3 for Sherlock; X, Y, Z = 13, −87, 7 for Summer).

**Fig 7.**
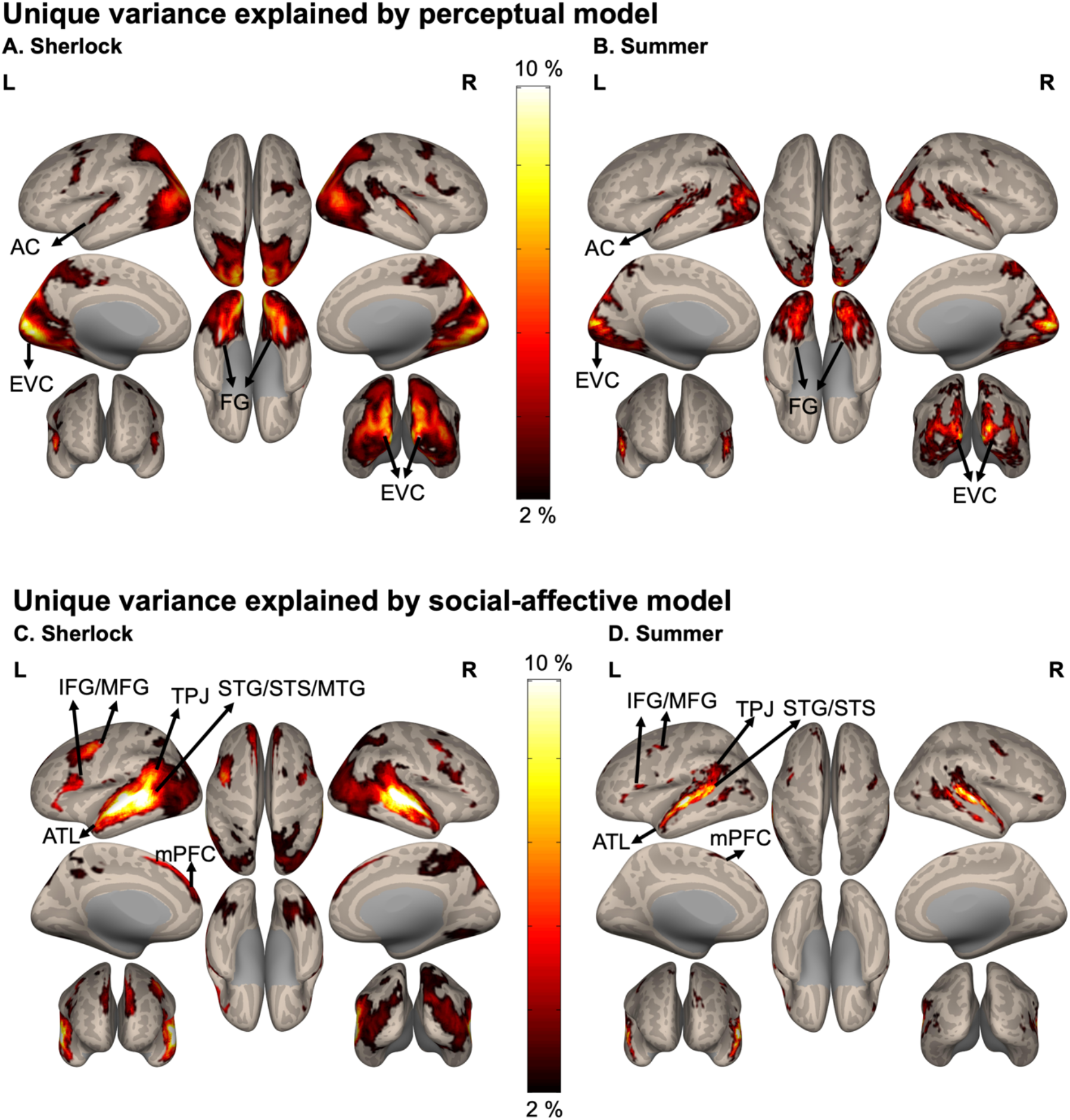
The portion of unique variance, expressed as percentages, explained by the perceptual (A, B) and social-affective model (C, D) in the Sherlock and Summer data. Color bar indicates percentage of unique variance explained. The perceptual model explained unique variance of neural responses in the visual and auditory cortices. In contrast, the social-affective model explained the unique variance of social brain responses, including the STS, TPJ, ATL, and mPFC activations. Group averaged variance partitioning maps are thresholded at P _FDR_ < 0.05, the minimum cluster size > 10 voxels, and the portion of explained unique variance > 2%. AC = auditory cortex, EVC = early visual cortex, FG = fusiform gyrus, IFG = inferior frontal gyrus, MFG = middle frontal gyrus, STG = superior temporal gyrus, STS = superior temporal sulcus, MTG = middle temporal gyrus, TPJ = temporoparietal junction, ATL = anterior temporal lobe, mPFC = medial prefrontal cortex. For labeling methods, see Variance Partitioning in Materials and Methods.

On the other hand, the social-affective model uniquely predicted the voxel-wise responses in high-level social cognitive regions, including the STS, TPJ, anterior temporal lobe (ATL), and mPFC in both studies (Fig 7C-D). The largest portion of unique variance explained was in the left STS at the same or immediately adjacent MNI coordinates where the social-affective model shows the highest performance. Note that a substantial portion of the shared variance explained by both models was observed in the temporal lobe and occipital lobe (data not shown). The fact that perceptual and social-affective features covary may explain this finding.

### The presence of a social interaction uniquely predicts brain response in the STS

Finally, we sought to identify whether two major, independent (though often correlated) social features, social interactions and theory of mind, independently predict brain activity during natural movie viewing. For the Sherlock data, the presence versus absence of social interactions uniquely predicted brain responses throughout the temporal lobe and to some extent in precuneus and inferior parietal sulcus (IPS), with the bilateral STS showing the strongest selectivity (Fig 8A). The largest portion of unique contribution was observed in the left STS at the same MNI coordinates, where the social-affective model showed the highest performance. The Summer fMRI data showed similar results, although the predicted brain areas were confined to right STS and bilateral precuneus, with the right STS (X, Y, Z = 52, −42, 7) showing the strongest selectivity (Fig 8B). This supports and extends the results of our feature preference mapping (Fig 6), highlighting the role of social interaction processing in the STS, independent of all other features, including spoken language.

**Fig 8.**
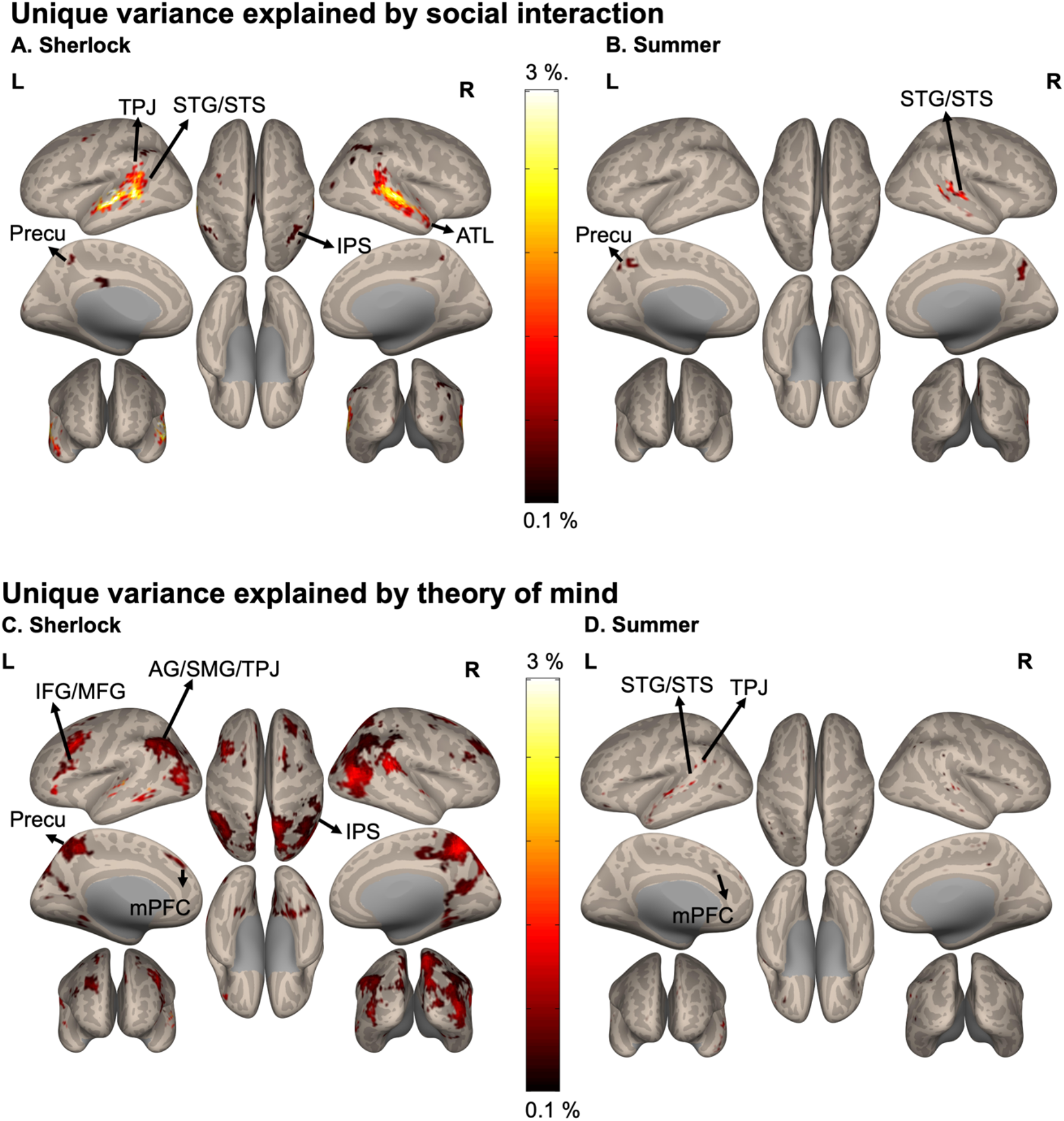
The amount of the unique variance explained by the social interaction (A, B) and theory of mind feature (C, D) in the Sherlock and Summer studies. Color bar indicates percentage of unique variance explained. The social interaction feature explained unique variance in neural responses in the STS and precuneus. The theory of mind feature explained the unique variance of neural responses in the theory of mind network, such as TPJ, precuneus, and mPFC. Group averaged variance partitioning maps are thresholded at P _FDR_ < 0.05 and the minimum cluster size > 10 voxels (except for the panel D, where the minimum cluster size > 1 voxel). STG = superior temporal gyrus, STS = superior temporal sulcus, MFG = middle frontal gyrus, ATL = anterior temporal lobe, mPFC = medial prefrontal cortex, TPJ = temporoparietal junction, AG = angular gyrus, SMG = supramarginal gyrus, IPS = intraparietal sulcus, Precu = precuneus

In the Sherlock data the theory of mind feature uniquely predicted neural responses of other social brain regions, mainly located in the theory of mind network (Dufour et al., 2013) – the precuneus, mPFC, and TPJ (Fig 8C), with the precuneus showing the strongest selectivity (X, Y, Z = 6, −60, 51). However, this distinct contribution of the theory of mind feature, observed in the Sherlock data, was not replicated in the Summer fMRI data. Only a few voxels, fewer than 10 voxels in each cluster, in STS, TPJ, and mPFC showed selectivity to the theory of mind feature (Fig 8D). Nonetheless, uncorrected variance partitioning results (i.e., P _uncorrected_ < 0.05) included the same brain regions reported in Sherlock, such as TPJ, precuneus, and mPFC, and were largely non-overlapping with the unique social interaction voxels. In addition, no voxel showed the shared variance explained by both social interaction and theory of mind in Sherlock data. We did not run this analysis in Summer data considering the weak results found in the theory of mind feature. Overall, the results show distinct functional and anatomical divisions between social interaction and theory of mind processing in natural movies. In particular, the STS shows strong, unique selectivity to scenes with social interactions, and the precuneus, mPFC, and TPJ show selectivity for scenes where characters infer others’ thoughts and emotions.

## Discussion

Here we investigated the brain mechanisms underlying naturalistic social interaction perception in a more ecologically valid and replicable context. Combining voxel-wise encoding and variance partitioning analyses (Fig 1), we identified the brain regions showing unique selectivity for general social-affective information (Fig 7C-D) and those particularly selective to social interactions (Fig 8A-B). We also demonstrated how auditory and visual features are encoded (Fig 7A-B), replicating prior findings (Kanwisher et al., 1997; Sunaert et al., 1999; Cohen et al., 2002; Hart et al., 2003; Brugge et al., 2009) in a natural setting. Importantly, our findings were replicated across both sets of movie data that came from different genres, subjects, and labs.

### Social-affective information processing during natural viewing

Movie viewing paradigms have recently been highlighted as essential tools in cognitive neuroscience due to their richness and comparable complexity to the real world (Sonkusare et al., 2019; Redcay and Moraczewski, 2020). Most prior social neuroscience studies with movies have used reverse correlation analyses to investigate phenomena such as theory of mind (Richardson et al., 2018) and social interaction perception (Wagner et al., 2016). These methods present a major advantage in that they do not require extensive movie labeling, but are prone to reverse inference errors as the event labeling happens post-hoc. This is particularly challenging in movies where the distribution of social information is often imbalanced (see Table 1). In the current study we densely labeled movies, using a combination of automatic and human annotations, and used the extracted movie features to perform cross-validated voxel-wise encoding analysis, which is less prone to reverse inference errors (Redcay and Moraczewski, 2020).

Using these methods, we found several brain regions that coded for social-affective information – social interaction, an agent speaking, mentalizing, perceived valence, and arousal – independent of co-varying perceptual information (Fig 7C-D). Specifically, we identified unique variance explained in brain regions previously attributed to social perception (STS) (Hooker et al., 2003; Vangeneugden et al., 2014; Lee Masson et al., 2018, 2020a; Pegado et al., 2018), theory of mind (TPJ, mPFC, ATL) (see the review in (Schurz et al., 2020), and action observation (inferior frontal gyrus (IFG)) (Carr et al., 2003; Centelles et al., 2011).

Surprisingly, the preferred feature across most of the brain, including many of the above social-affective regions, was the fifth layer of a DNN pre-trained on an object recognition task (Fig 6). This result is hard to interpret as most prior neuroimaging studies using DNNs have focused on their match to voxel responses in the ventral visual stream (Khaligh-Razavi and Kriegeskorte, 2014; Güçlü and van Gerven, 2015; Bonner and Epstein, 2018; Zeman et al., 2020). One prior study found that object-trained DNNs predict voxel-wise responses in the TPJ (Wen et al., 2018), concluding that later DNN layers might capture high-level semantic information in naturalistic visual scenes. Further investigation is needed to better understand the substantial contribution of late-stage DNN layers in explaining social brain activity. Intriguingly, the most notable exception to DNN preference was preference for social interactions in the STS. It is difficult however, to rule out the effect of spoken language processing (Deen et al., 2015; Wilson et al., 2018) on STS activity with this analysis alone. Although we tried to capture this information with our auditory and spoken language features, it is likely these do not capture all of the rich semantic content present in the movie. As the semantic network largely overlaps with brain areas observed in the current study (Huth et al., 2016; de Heer et al., 2017), future investigations should use language models to understand how semantic content overlaps with different types of social perception. With this limitation in mind, we turned to variance partitioning, to identify the unique contribution of social interactions to brain responses.

### Social interaction perception during natural viewing

In both studies, the right STS and, to a lesser extent, the precuneus showed unique selectivity for naturalistic social interactions (Fig 8A-B), independent of all other covarying features. These results provide the first evidence that unique variance in STS is explained by the presence/absence of social interactions during natural viewing. Although we replicated our findings across two movie datasets, we observed a slight discrepancy with respect to the extent of STS activity and the lateralization. This discrepancy has also been observed in other studies (e.g., (Isik et al., 2017; Lee Masson et al., 2018; Walbrin et al., 2018) found right lateralization, while others did not (Lahnakoski et al., 2012; Walbrin and Koldewyn, 2019; Walbrin et al., 2020)). Our findings highlight that although STS is known as a hub for general social processing, social interaction is a critical feature that uniquely contributes to STS responses.

Unique variance in the precuneus was also explained by social interaction, but to a lesser degree than STS. The precuneus has been implicated in a wide range of cognitive processes, including false belief tasks (Saxe and Kanwisher, 2003; Jacoby et al., 2016), social trait judgment tasks (Farrer and Frith, 2002; Iacoboni et al., 2004), and while viewing social interactions (Iacoboni et al., 2004; Lahnakoski et al., 2012). A possible interpretation of our findings is that the STS and precuneus work in concert to spontaneously extract socially relevant information about others. This notion is supported by previous studies showing increased precuneus’s connectivity with other social brain areas during the observation of dyadic interaction compared to human-object manipulation (Lee Masson et al., 2020b) and social evaluation task on others’ faces (McCormick et al., 2018).

A prior study found increased response to social interaction scenes of a movie in mPFC, concluding that a viewer may spontaneously infer movie characters’ thoughts and intentions (Wagner et al., 2016). Contrary to that finding, we did not observe mPFC involvement specific to social interaction perception after controlling for the effects of perceptual and social features, including speaking and theory of mind. Instead, mPFC was selectively tuned to more general social-affective features (Fig 7C-D) and theory of mind (Fig 8C-D). Notably, in Wagner and colleagues’ study, movie scenes that evoked strong mPFC responses were speaking scenes identified with reverse correlation analysis, and they did not consider theory of mind features in their analysis. Given that social interaction scenes genuinely invite theory of mind (Roeyers et al., 2001; Dziobek et al., 2006; Grainger et al., 2019), it may make more sense to compare their findings to our findings on general social-affective features that include speaking, social interaction, and theory of mind. Our variance partitioning results show for the first time that social interaction perception is separable from theory of mind processing, even in natural conditions.

### Theory of Mind during natural viewing

Recent results have shown increased responses in the theory of mind network, including TPJ, precuneus, and mPFC, during movie viewing (Jacoby et al., 2016; Richardson et al., 2018). While we largely replicated these results in the Sherlock data, we did not observe the same in the Summer data. This discrepancy may be related to the substantially lower signal-to-noise ratio in the Summer movie data (maximum ISC value 0.73 vs. 0.56, see inter-subject brain correlation in Methods). Another important distinction is how we annotated the theory of mind feature. Unlike previous studies (Jacoby et al., 2016; Richardson et al., 2018), our theory of mind annotation was labeled in advance based on whether the scene contained a person speaking about others’ mental states. We believe this criterion to be more objective than trying to guess whether the subjects were engaged in mentalization. However, it may pose issues for certain movie scenes. For example, 500 Days of Summer (unlike Sherlock) contains many scenes with a narrator describing characters’ mental states in voice-over monologues. We did not distinguish between scenes with the narrator and examples of mentalization that happened more naturally in the course of the movie.

### Future directions

The methods and findings presented in this study open the door for many avenues of naturalistic social neuroscience research. Our results suggest that, even in a real-world setting, social interactions are processed in a manner that is distinct from other perceptual and social features. In both simple visual displays (Su et al., 2016) and natural images (Skripkauskaite et al., 2021), there is an attentional bias for social interactions. Why do social interactions capture our attention and how do we use them to reason about interacting individuals? Is there a computational advantage to selectively processing social interactions?

Finally, our method of isolating brain areas selective to naturalistic social interaction in MRI may be particularly advantageous for studies of typical and atypical development, including autism. Many high functioning individuals with autism look identical to neurotypical adults in simple social psychology and neuroscience tasks despite differences in their real-world social abilities (Scheeren et al., 2013; Moessnang et al., 2020). The methods outlined here may help us close the gap between simple lab-based tasks and the real world.

## Acknowledgements

We thank Janice Chen, Sarah Aliko, Jongwan Kim and their colleagues for data and annotations used in this study. We would like to thank J. Brendan Ritchie and Alon Hafri for valuable comments. This work was supported with funds from The Clare Boothe Luce Program for Women in STEM.

